# A20 is an immune tolerance factor that can determine islet transplant outcomes

**DOI:** 10.1101/770834

**Authors:** Nathan W. Zammit, Stacey N. Walters, Karen L. Seeberger, Gregory S. Korbutt, Shane T. Grey

## Abstract

Islet transplantation can restore lost glycemic control in type 1 diabetes subjects, but is restricted in its clinical application by limiting supplies of islets and the need for heavy immune suppression to prevent rejection. *TNFAIP3*, encoding the ubiquitin editing enzyme A20, regulates the activation of immune cells by raising NF-κB signalling thresholds. Here we show that increasing A20 expression in allogeneic islet grafts resulted in permanent survival for ~45 % of recipients, and >80% survival when combined with subtherapeutic rapamycin. Allograft survival was dependent upon regulatory T cells, was antigen-specific and grafts showed reduced expression of inflammatory factors, but increased TGFβ and IL-10. By analysing islets expressing an A20 coding mutation (I325N) that cripples A20’s OTU ubiquitin editing domain, we found that A20 regulates intra-graft RIPK1 levels to modulate NF-κB signalling. Transplantation of I325N islets resulted in increased NF-κB signalling, graft hyper-inflammation and acute allograft rejection. Neonatal porcine islets (NPI) represent a clinical alternative islet source but are readily rejected. However, forced A20 expression reduced NPI inflammation and increased their function after transplantation. Therapeutic administration of A20 raises NF-κB signalling thresholds and promotes islet allogeneic survival. Clinically this would allow for reduced immunosuppression supporting the use of alternate islet sources.

## Introduction

Type 1 diabetes (T1D) is an autoimmune condition marked by loss of glycemic control caused by immune-mediated destruction of insulin producing beta cells that reside within the pancreatic islets of Langerhans (1). Replacement of lost beta cells by adult islet allogeneic transplantation restores glycemic control, providing fine-tuned release of insulin in response to blood glucose in real-time, something not yet achievable by manual or automatic injection of insulin or its analogues (2–4). Islet transplantation reduces exogenous insulin requirements and reverses hypo-glycemic unawareness, a life-threatening complication of T1D (3, 5, 6). Although highly successful, the need for robust suppression of host immunity to avoid rejection precludes its indication for pediatric T1D patients, restricting the broader application of islet transplantation to adults with life threatening hypoglycemic unawareness (2, 7).

Islet transplantation is further restricted by the scarcity and fragility of islets. Frequently patients require multiple islet infusions extracted from multiple pancreata to achieve clinical outcomes of insulin independence and reversal of hypo-glycemic unawareness (4, 6, 8). Further to this, most islet transplant recipients show relatively poor long-term outcomes compared to solid organ transplant recipients, requiring a return to insulin injections within a few years post-islet transplant (2, 4). Evidence suggests the underlying mechanisms leading to reduced islet allograft survival are unique to islet transplantation and include recurrent islet autoimmunity, sensitivity of islets to the intra-portal transplant site and islet-toxicity of immunosuppressive drugs, as well as factors present in solid organ transplantation such as chronic allograft rejection (2). These factors are likely exacerbated by autologous islet inflammation induced via the isolation process and *ex vivo* culture that may hasten graft failure and increase their immunogenicity after transplant (9–11). Strategies that reduce islet fragility and inflammation could preserve islet graft-mass and improve post-transplant function potentially reducing the reliance on heavy immunosuppression and widening the eligibility criteria for an islet transplant (10).

*TNFAIP3*, encoding the ubiquitin editing protein A20, is a master regulator of NF-κB signalling. A20 through its ovarian tumor (OTU) and zinc finger 4 domain modifies ubiquitin chains on key intracellular inflammatory signalling mediators, primarily RIP1 and TRAF6 (12, 13) that lie downstream of inflammatory and danger sensing receptors of the TNF receptor family, including TNFR1, IL-1R and TLRs. In hematopoietic cells A20 functions as a negative regulator of immuno-stimulatory factors and thus governs the threshold for immune activation. Reduced expression of A20 in dendritic cells leads to increased expression of costimulatory molecules and an enhanced ability to activate CD8^+^ and CD4^+^ T cells during an immune response (14–16). Further, deletion of A20 in B cells, macrophages and granulocytes results in cell intrinsic hyper-activation and spontaneous inflammatory disease in mice (17–20). In human subjects A20 haploinsufficiency is associated with increased serum cytokines, higher frequencies of TH17 cells and autoimmune disease (21, 22). Thus, by regulating NF-κB activation, A20 sets the threshold for the generation of a productive immune response. Here we investigated the impact of changed A20 expression levels in islet allografts on immune-stimulatory thresholds and islet allograft survival.

## Results

### Forced expression of A20 allows permanent islet allograft survival without needing immunosuppression

To test the impact of A20 on tissue tolerance, we transduced primary islets with an adenoviral vector (rAd.A20) to force A20 expression to high levels (Figure 1A and Supplemental Figure 1), which resulted in a robust suppression of the normal pro-inflammatory gene response following TNF and IL1β stimulation (Figure 1, B and C). Further analysis in beta cell lines showed that increasing A20 levels suppressed TNF-induced NF-κB and JNK signalling pathways, inhibited activation of a NF-κB and a AP-1 reporter, and suppressed expression of pro-inflammatory factors associated with allograft rejection (Supplemental Figure 2) (9). Next, primary islets from BALB/c (H2^d^) donor mice transduced with rAd.A20 were transplanted into diabetic C57Bl/6 (H2^b^) allogeneic recipients. Adenoviral transduction did not affect islet graft function *in vivo* as demonstrated by the ability of both rAd.A20, control rAd.GFP (GFP-expressing) and non-infected (NI) grafts to rapidly restore euglycemia following transplantation (Figure 1D). Kaplan-meier survival analysis showed rapid rejection of control NI and rAd.GFP transduced islet grafts allografts (Figure 1E). In contrast, ~ half of mice receiving A20 expressing islets failed to reject their grafts and instead exhibited permanent (>200 days) allograft survival. Graft removal by survival nephrectomy for some recipients at post-operative day (POD) 100 disrupted glucose control, illustrating that A20-transduced surviving islet grafts were both functional and responsible for euglycemia (Figure 1F). Long-term surviving A20-transduced grafts were characterised by normal islet architecture, robust insulin production and distinct pockets of mononuclear cells within the graft microenvironment (Figure 1G). Enhanced graft morphology was also evident for A20-expressing grafts at POD 10 (Figure 1H). Thus, forced expression of A20 allows permanent and functional survival of an islet allograft without needing immunosuppressive drugs.

**Figure 1.**
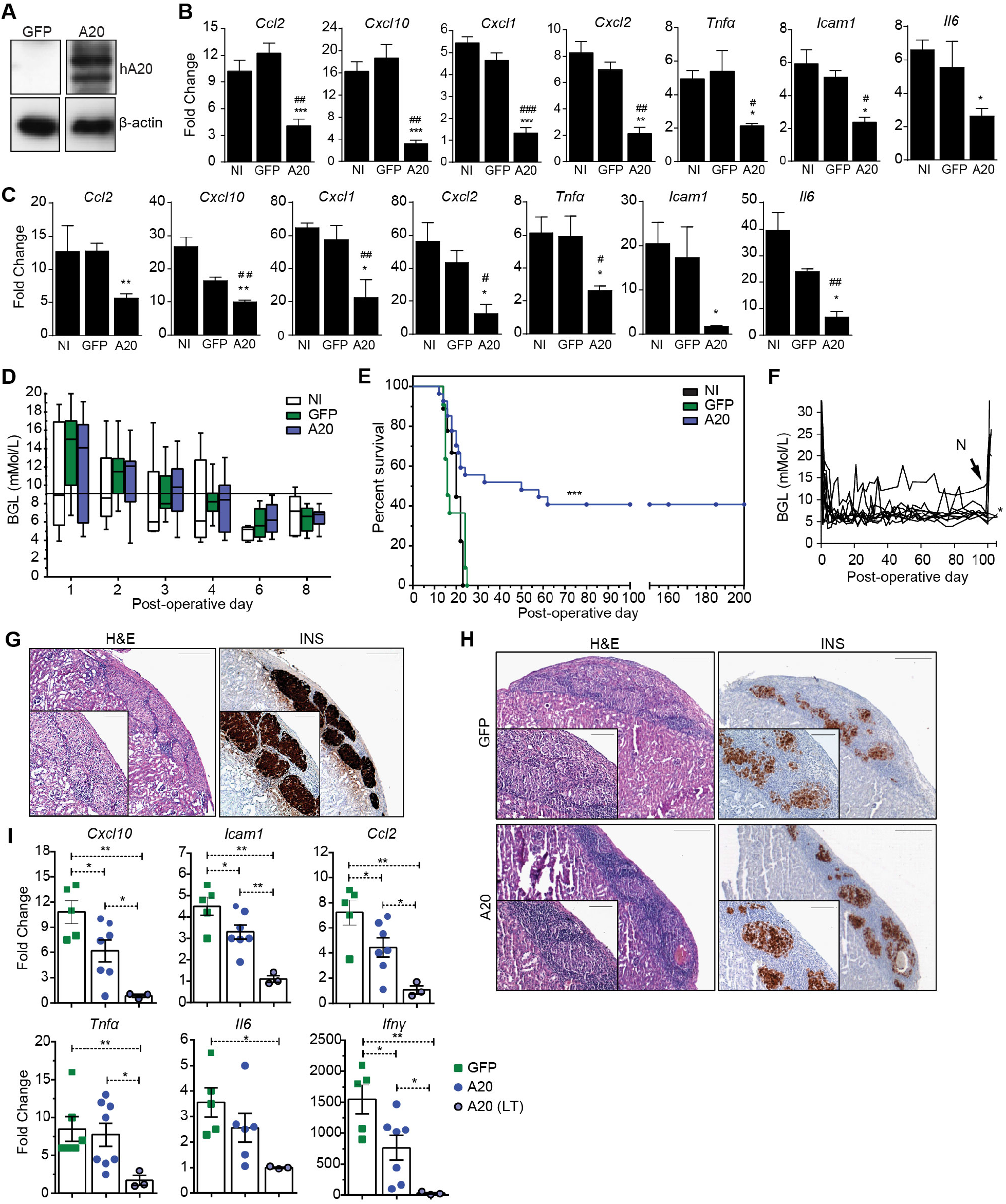
Improved survival characteristics of an A20-expressing islet allograft. Primary islets transduced with adenoviral constructs encoding for GFP, A20 or left non-infected (NI). Following 24 h islet grafts were **(A)** lysed and A20 protein levels assessed by immunoblot, or **(B, C)** treated with 200 U/ml of TNFα (B) or IL-1β (C) for 4 h and mRNA levels of inflammatory factors assessed by RTPCR. (* = A20 versus NI; # = A20 versus GFP). **(D-E)** GFP (*n* = 9; p=0.16), A20 (*n* = 27; *p* = 0.002) and NI islet grafts (*n* = 11) were transplanted under the kidney capsule of allogeneic C57BL/6 mice and (D) Blood glucose levels (BGL) and (E) survival of islet grafts monitored for the indicated days. **(F)** Nephrectomies (N) were conducted at post-operative day (POD) 100 for a portion of A20-expressing long term serving islet grafts. **(G, H)** Hematoxylin & Eosin staining (H&E) or insulin labelling (INS) of long-term (>100 days) surviving grafts (G) or of GFP or A20 expressing grafts at POD10 (H). Scale bar = 200 μm [low-power field] 100 μm [high-power field]. **(I)** GFP (closed square) or A20 (closed circle) expressing grafts harvested at POD10, as well as A20-transduced long-term surviving grafts harvested at > POD100 (open circle) and subjected to qRT-PCR for mRNA levels of islet derived inflammatory factors and IFNγ. Error bars ± SEM and *P* values represent Student’s *t*-test unless otherwise stated, * *P* < 0.05; \*\**P* < 0.01.

### Immune features of A20-induced islet allograft survival

We investigated the immunological mechanism for long-term survival of A20-expressing allografts. After 150 days post transplantation, splenic T cells were harvested from mice with A20-expressing BALB/c (H2^d^) islet grafts and were adoptively transferred to RAG^−/−^ mice previously transplanted with a BALB/c (H2^d^) islet allograft. Control groups received splenic T cells harvested from C57/BL6 mice (Figure 2A). In this situation the majority of RAG^−/−^ mice receiving T cells taken from mice with surviving A20-expressing grafts took longer to reject their islet grafts, and the majority permanently accepted the allograft, compared to RAG−/− mice receiving T cells from naïve C57/BL6 mice (Figure 2B). Thus, A20-induced islet allograft acceptance is T cell dependent. To determine whether graft acceptance was due to T cell anergy, deletion or regulation, we repeated the above experiment but this time only transferred FACS-purified CD4+ CD25-effector T cells from mice harbouring long-term surviving A20-expressing or control non-infected islet grafts. These T cell preparations lacked CD4+ CD25+ T cells with regulatory potential (23, 24). In this experiment all of the recipient mice rejected the second BALB/c allograft regardless of whether they received effector T cells from mice with A20-expressing grafts or control grafts (Figure 2B). This indicated to us that A20-expression engendered T cell dependent tolerance. To test if graft acceptance was specific to the BALB/c (H2^d^) alloantigen, we established another cohort of long-term surviving A20-expressing islet graft recipient mice to repeat the above experiment. However, in this case the T cells from mice harbouring A20-expressing long-term surviving grafts were adoptively transferred into RAG^−/−^ mice pre-transplanted with a MHC-disparate graft from a different (H2^k^) donor strain (Figure 2C). Subsequently, in all cases the H2^k^ MHC-mismatched grafts were rapidly rejected. We conclude from these experiments that A20-expression does not delete effector T cells or impair their function, or induce T cell anergy, but rather results in T cell dependent and antigen specific immune regulation towards an islet allograft.

**Figure 2.**
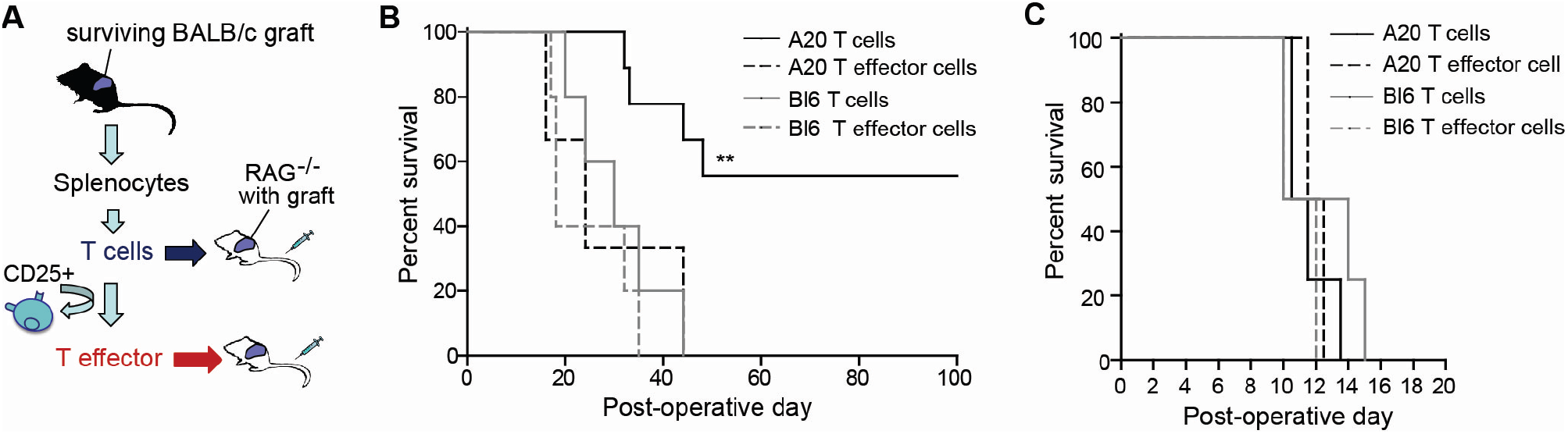
A20-induced islet allograft survival is T cell dependent and is antigen specific. **(A)** Experimental strategy and **(B)** Kaplan-Meier survival data of RAG^−/−^ mice pre-transplanted with a BALB/c graft and adoptively transferred with T cells or CD25 depleted T effector cells, from mice harbouring long-term surviving grafts (A20 T cells, *n* = 9; A20 T effector cells., *n* = 3) or control C57BL/6 mice (B6 T cells, *n* = 5; B6 T effector cells *n* = 5). **(C)** Kaplan-Meier survival data of RAG^−/−^ mice pre-transplanted with a third-party CBA (H2^k^) graft and adoptively transferred with T cells or CD25 depleted T effector cells from mice harbouring long-term surviving grafts (A20 T cell, *n* = 4; A20 T effector cell., *n* = 2) or control C57BL/6 mice (Bl6 T cell, *n* = 4; B6 T effector cell, *n* = 2). Log rank test used for significance \*\**P*<0.01.

### Forced expression of A20 engenders Foxp3^+^ T cells

Both A20- and GFP-expressing islet allografts were infiltrated with FOXP3^+^ cells at POD 10 after transplantation (Figure 3A) but the number of FOXP3^+^ cells within the GFP-graft microenvironment fell during the time period when grafts were being rejected between POD15-25. In contrast, A20-expressing grafts maintained high numbers of FOXP3^+^ cells (Figure 3, B and D). Also, long-term surviving A20-expressing grafts (>100 days) showed prominent infiltration of FOXP3^+^ cells congregated within the peri-graft space appearing to surround each individual islet (Figure 3, C and D). The presence of FOXP3^+^ cells always correlated with improved islet graft architecture, whereas the immunopathology of GFP-expressing grafts at POD10 revealed reduced number of FOXP3^+^ cells with fragmented, less defined islet architecture and patchy insulin labelling compared to the well-preserved islet architecture of A20-expressing grafts at POD10 and POD100 (Figure 3A-C).

**Figure 3.**
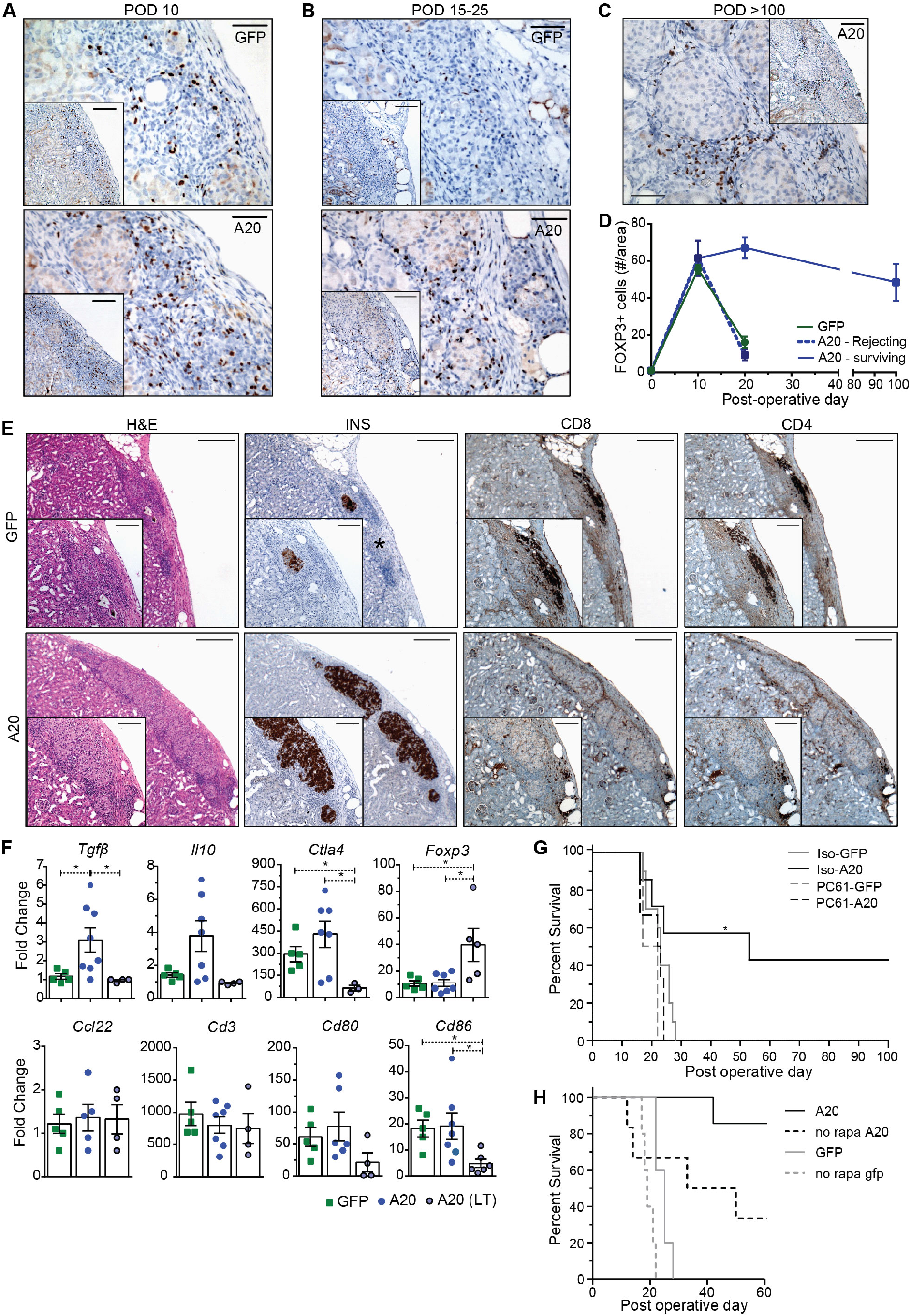
Long-term surviving grafts have graft infiltrating FOXP3^+^ cells. **(A)** Representative sections of FOXP3 stained GFP or A20 transduced allografts at post-operative day (POD)10, **(B)** POD 15-25 (GFP grafts taken before rejection) and, **(C)** POD>100. **(D)** Quantification of FOXP3^+^ cells. Scale bar = 100 μm (low-power field), 50 μm (high-power field). **(E)** Representative Hematoxylin & Eosin staining (H&E), insulin (INS), CD8 or CD4 labelling of GFP or A20 expressing grafts at POD 18 (Scale bar = 200 μm [low-power field] 100 μm [high-power field]; * denotes islet no-longer producing insulin). **(F)** GFP or A20 expressing grafts harvested at POD10, as well as, A20-expressing long-term surviving grafts harvested at >POD100 and subjected to RT-PCR for known immune regulatory factors. **(G)** Survival of BALB/c islet grafts expressing GFP or A20 when transplanted into allogeneic C57/BL6 recipients given repeated doses of αCD25 PC61 (PC61-GFP, *n* = 2; PC61-A20, *n* = 6) or an isotype-control (Iso-GFP, *n* = 10; Iso-A20, *n* = 7). **(H)** Survival data for C57BL/6 mice receiving allogeneic BALB/c islet grafts transduced with GFP or A20 (MOI = 10:1). Recipient mice were either given a sub-therapeutic dose of Rapamycin (0.1 mg/kg at day of transplantation and every day thereafter for 7 days (A20 *n*=7, GFP *n*=5), or no Rapamycin (A20 *n*=7, GFP *n*=5). Error bars ± SEM and *P* values represent Student’s *t*-test unless otherwise stated, * *P* < 0.05.

The increased number of Foxp3^+^ cells in A20-expressing grafts was associated with reduced levels of pro-inflammatory mRNA in the graft site at POD10 that associate with islet graft rejection (9, 25) as compared to expression levels observed for GFP-expressing grafts (Figure 3F). These same mRNAs were also suppressed in long-term surviving A20-expressing grafts (Figure 3F). Further to these data we found a noticeable reduction in the frequency of CD8^+^ and CD4^+^ positive cells within the A20-expressing grafts (Figure 3E). However, A20 expressing grafts exhibited elevated levels of *Tgfβ* and a trend to increased levels of *Il10* and *Ctla4* within the graft microenvironment (Figure 3, F). However, *Ccl22*, a chemokine that attracts Tregs (26), was not found to be differentially expressed between groups. There was also no overall change in the level of dendritic cell activation markers *Cd80* or *Cd86* between A20- or GFP-expressing grafts at POD 10 (Figure 3, F). Within long-term surviving grafts (>100 days) FOXP3^+^ cells and high levels of *Foxp3* mRNA was readily detected (Figure 3, C and F). Therefore A20 alters the pro-inflammatory millieu and engenders regulatory T cells within the graft microenvironment.

To further investigate the role of regulatory T cells in A20-induced tolerance we treated diabetic recipient mice with the αCD25 mAb PC61, which depletes CD25^+^ FOXP3^+^ regulatory T cells (27) (Supplemental Figure 3), by preventing CD25 binding to interleukin 2 (28, 29). In this experiment, all of the mice receiving A20-transduced grafts and treated with PC61 mAb rapidly rejected their grafts with similar rejection times to those observed for control GFP-expressing grafts (Figure 3G). In contrast, 40% of recipients of A20 expressing grafts injected with an isotype control antibody exhibited long-term survival (Figure 3G). We conclude that graft intrinsic expression of A20 can promote regulatory T cell dependent tolerance to an MHC mismatched islet allograft.

Rapamycin (also known as sirolimus) inhibits mammalian target of rapamycin (mTOR) and is used clinically as an immunosuppressant to dampen T cell responses in transplantation (4, 6). Preclinical and clinical studies also indicate that rapamycin promotes FOXP3^+^ regulatory T cells (30–33), therefore we investigated whether graft intrinsic expression of A20 would synergise with the tolerance promoting properties of rapamycin. For the experiment diabetic C57BL/6 recipients received A20-expressing H-2^d^ BALB/c islet allografts, as well as seven daily injections of a limiting subtherapeutic dose of rapamycin (25) starting on the day of transplantation (Figure 3H). All control GFP-expressing grafts, and those treated with rapamycin alone, were rapidly rejected, whereas mice receiving A20-expressing grafts and treated with a subtherapeutic dose of rapamycin showed superior graft survival compared to grafts transduced with A20 alone (Figure 3H). These latter data highlight the translational potential of A20 to synergise with clinical approaches to promote significant improvements in islet allograft outcomes.

### A20 promotes tissue tolerance by regulating RIPK1

As increasing intra-graft A20 levels can promote allograft tolerance by increasing the threshold for NF-κB activation we investigated whether A20 reduction would have the reverse effect and promote inflammation with more aggressive allograft rejection. To test this we utilised an ENU-mutagenesis generated mouse line harbouring a germline A20 loss of function mutation (34). This coding mutation lies within A20’s functional OTU ubiquitin editing domain (12) and substitutes a conserved isoleucine at amino acid position 325 for an asparagine (I325N). When transiently expressed in pancreatic beta cell lines the I325N mutation cripples A20’s ability to inhibit TNF-induced NF-κB and JNK reporter activation (Figure 4A-C), and when overexpressed in wild-type mouse islets the I325N A20 variant shows a reduced ability to inhibit TNF-induced inflammatory gene expression as compared to islet expressing wild-type A20 (Figure 4D). A20 regulates inflammatory signalling by terminating RIPK1 activation via cleavage of K63 ubiquitin chains with its OTU domain and targeting substrates for K48-mediated proteosomal degradation via its zinc finger 4 ubiquitin ligase domain (12). The I325N mutation does not alter A20 protein stability, nor A20’s capacity to interact with key substrates RIP1K1 or NEMO in beta cells (Supplemental Figure 4). Rather, the I325N mutation resulted in increased accumulation of hyper-polyubiquitinated isoforms of RIPK1 (Supplemental Figure 4A) and increased RIPK1 levels in TNF simulated cells (Supplemental Figure 4, A and C), consistent with a reduction in A20’s ubiquitin editing function (12). Further to this, islets isolated from I325N mice exhibited increased TNF-induced proinflammatory signalling and gene expression compared to wild type islets (Figure 4, E and F). When transplanted into diabetic allogeneic recipients I325N islets showed accelerated rejection, and a hyper-inflammatory graft microenvironment with reduced expression of factors associated with immune regulation including *tgfb*, *l10* and *ctla4* (Figure 4G-I). Thus, A20 is necessary to control islet homeostasis in response to inflammatory triggers. In the specific context of islet transplantation changing A20 levels can function as an immune modulatory control switch that dictates transplant outcomes.

**Figure 4.**
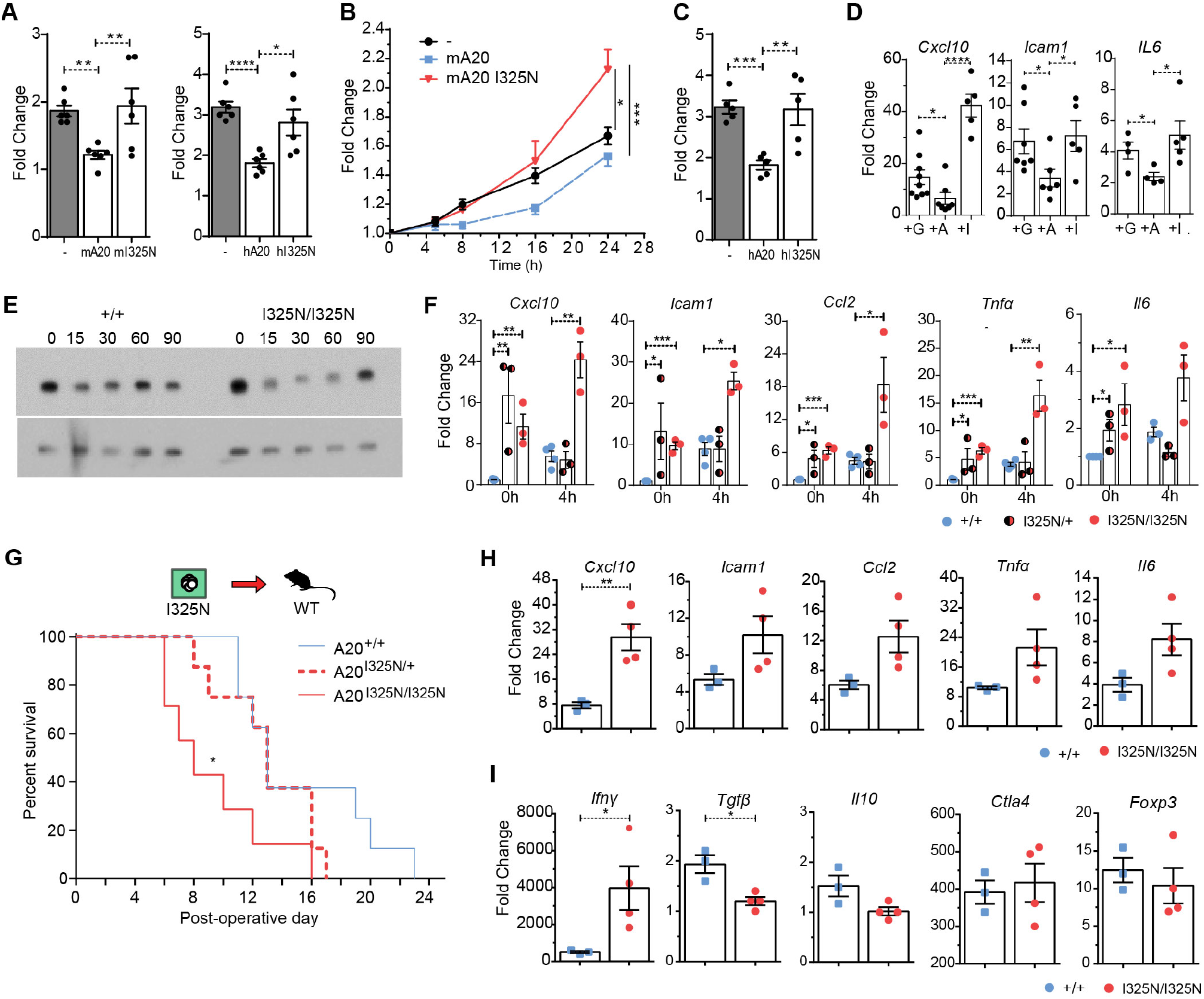
Reduced A20 function leads to rapid islet allograft rejection. **(A-C)** βTC3 cells co-transfected with an NF-κB.luciferase reporter (A), or an AP-1 luciferase reporter (B, C) and a CMV.βgal expression construct with or without PCDNA3.1 encoding murine (m) or human (h) reference A20, or A20 with an I325N coding variant. Cells were stimulated with 200 U/ml TNFα for 8 h (A, C), or for 5, 8, 16 and 24 h (B), or left untreated. Data represents fold change of stimulated versus non-stimulated. **(D)**Wild-type BALB/c islets were isolated and transduced with rAd.*GFP* (+G), rAd.*TNFAIP3* (+A) or rAd.*TNFAIP3*^I325N^ (+I), incubated overnight and stimulated with 200 U/ml TNFα for 0 or 4 h and mRNA levels for proinflammatory factors assessed by RT-PCR. **(E)** Immunoblot for IκBα and β-actin (loading control) of whole lysates from islets isolated from *Tnfaip3*^I325N/I325N^, or *Tnfaip3*^+/+^ mice and treated with 200 U/ml TNF for the indicated times. **(F)**Expression of TNF-induced genes in islets from *Tnfaip3*^+/+^, *Tnfaip3*^I325N/+^, *Tnfaip3*^I325N/I325N^ mice. Data represents 3 independent islet preparations with 3 biological replicates. **(G)**Kaplan-Meier survival data for *Tnfaip3*^I325N/I325N^ (*n* = 8; mean survival time [MST] = 8; p = 0.014), *Tnfaip3*^I325N/+^ (n = 8; MST = 13) or *Tnfaip3*^+/+^ (*n* = 8; MST = 13) islet grafts (H2^b^) transplanted under the kidney capsule of streptozotocin-induced diabetic CBA (H2^k^) mice (Log-rank test). **(H, I)** Islet grafts were excised at post-operative day 10 and subjected to RT-PCR for mRNA levels of known islet derived inflammatory factors (H) and non-islet derived factors (I). Non-transplanted overnight cultured isolated islets were used as base-line. Error bars represent SEM and Student’s *t*-test used for significance analysis unless otherwise stated, \**P*<0.05; \*\**P*<0.01; ***P<0.001.

### Forced expression of A20 improves the metabolic function of neonatal porcine islets post transplantation

In the clinical scenario adult human islets are transplanted into patients with T1D and hypoglycemic unawareness (5). The supply of suitable organ donors limits clinical islet transplantation and neonatal porcine islets (NPI) may be able to bridge that gap (35–38). We tested whether the protective and anti-inflammatory properties of A20 would transfer to NPI and thus demonstrate the potential for clinical translation. We transduced NPI with rAd.A20 to achieve high levels of A20 expression without evidence of toxicity, which blunted TNF-induced NF-κB and JNK signalling pathways, and suppressed expression of pro-inflammatory factors (Figure 5A-C; Supplemental Figure 5). Suppressed genes included tissue factor (TF) (Figure 5D), which plays a critical role in triggering posttransplant coagulation and enhances islet graft loss (39, 40). To test the beneficial efficacy of A20 *in vivo*, rAd.A20 transduced NPI were transplanted under the renal capsule of diabetic RAG^−/−^ mice and the time to normalisation of blood glucose was followed. A20-expressing NPI provided superior blood glucose control illustrated by a more rapid normalisation of posttransplant blood glucose and a normalised glucose tolerance (ipGTT) compared to mice receiving control NPI grafts (Figure 5, E and F).

**Figure 5.**
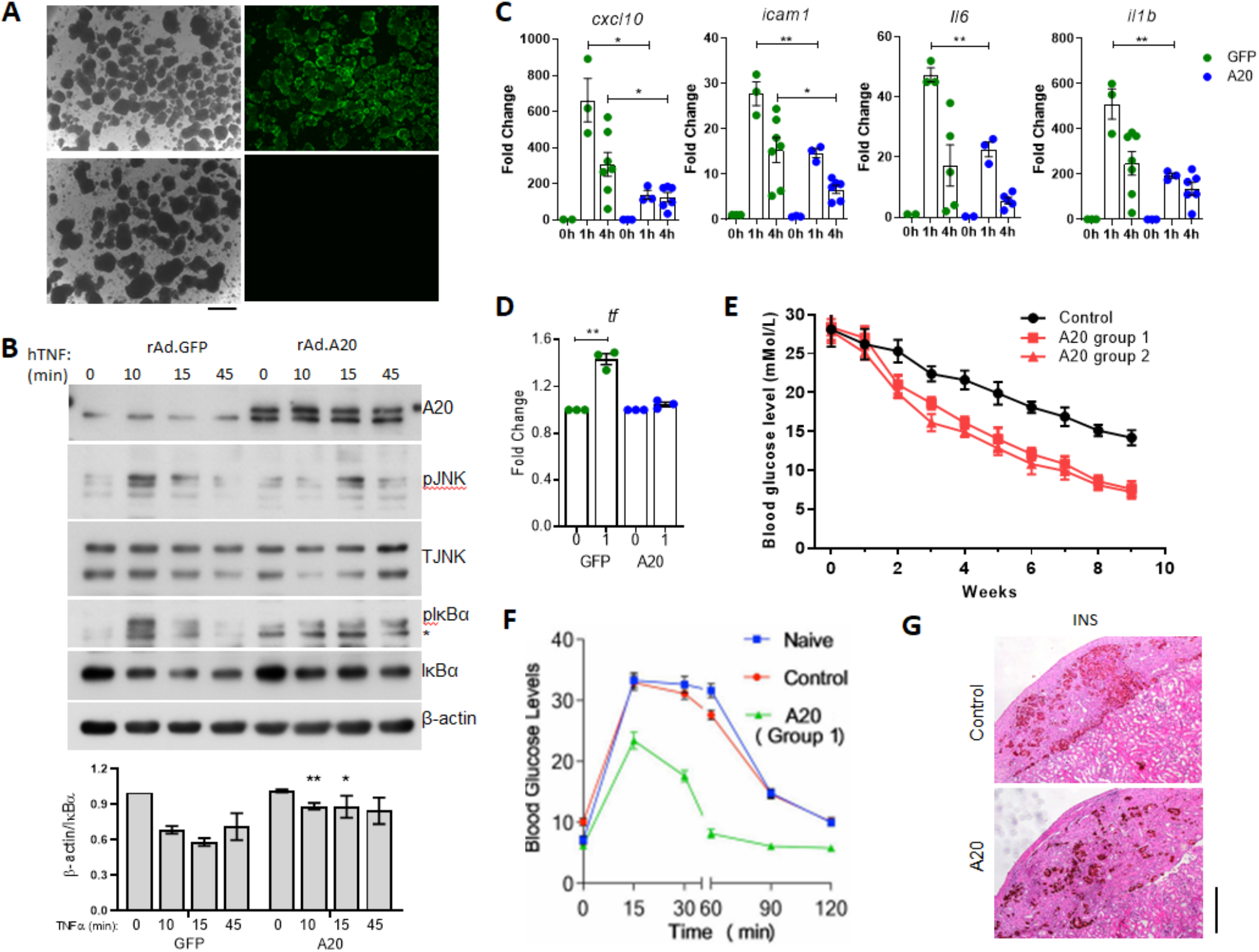
A20 expression in neonatal pig islets improves islet function following transplantation. **(A)** Micrograph image of neonatal porcine islet like clusters (NPI) transduced with an adenovirus encoding for GFP at a multiplicity of infection (MOI) of 10:1, or left untreated (scale – 200 μm). **(B)** Representative immunoblot of lysates from NPIs transduced with GFP or A20 at an MOI of 10:1 and stimulated with TNF for the indicated times. Bottom panel shows cumulative densitometry of IkBa levels from 4 independent experiments. **(C)** Transduced islets were also cultured overnight then with 200 U/ml TNF for indicated times and mRNA levels of inflammatory genes and **(D)** for tissue factor assessed. Data pooled from 3 independent experiments. **(E)**Average weekly blood glucose levels for diabetic RAG KO mouse cohorts (n=7) receiving 3000 NPI expressing either GFP or A20. **(F)** At 9 weeks post transplantation an intraperitoneal glucose tolerance test was performed, **(G)** and grafts collected for insulin (INS) staining. Scale = 200 μm.

## Discussion

Here we investigated the impact of changing A20 expression levels in islet allografts on immune-stimulatory thresholds and islet allograft survival. A potential role for A20 in transplantation was first indicated in a rodent heart xenotransplantation model where it was found that surviving hearts showed intra-graft A20 expression compared to rejecting hearts (41). Further studies showed that A20 reduces endothelial inflammation during xenotransplantation (42), the severity of graft arteriosclerosis (43) and improves liver graft function (44). Regarding pancreatic islets and beta cells, A20 expression is transcriptionally regulated in beta cells by NF-κB (45) but also involved in negative regulation of NF-κB activation (46). This suggests that manipulating A20 levels in islets would have a good safety profile but also have clinical potential as an approach for the suppression of otherwise deleterious NF-κB-dependent inflammatory genes (10). This is supported by studies showing A20 to be a potent inhibitor of NF-κB-mediated inflammation and cell death in pancreatic islets (47–49), and improving metabolic control of islets in a syngeneic transplant model when overexpressed (50). Here we demonstrate for the first time that islet-A20 dictates islet allograft fate. A20 overexpression engenders the accumulation of intra-graft antigen-specific regulatory T-cells that provide local tissue tolerance and islet graft survival. In contrast loss of A20 function accelerates rejection.

Our data fits with accumulating evidence in various tissues that show *TNFAIP3* to be a crucial gene for the maintenance of tolerance via its role in dampening responses to NF-κB-inducing inflammatory mediators. Indeed, A20 expression increases the threshold for immune cell activation, and reducing A20 in dendritic cells (14–16), T cells (51), B cells (17, 18), macrophages and granulocytes (19), results in their autosomal activation, with improved antigen presenting activity in the case of A20-deficient dendritic cells and spontaneous inflammatory disease in mice. In addition, A20 plays a central role to maintain intestinal tolerance, where a loss of A20 leads to aberrant responses to gut microbiota with intestinal inflammation (20, 52). The requirement of A20 for tissue homeostasis is highlighted by the fact that the A20 locus has been associated with many autoimmune and inflammatory disorders by GWAS (53). Further to this, A20 haploinsufficient human subjects present with increased TH17 cells and inflammatory disease (21, 22), whereas mice with A20 germline deficiency succumb to uncontrolled inflammation (54).

Here we show that manipulating A20 levels to achieve tissue tolerance can be directed to improve islet transplantation outcomes. Future studies will focus on generating clinical grade vectors for the delivery of A20. The continued maturation of gene delivery technology and gene editing techniques in pig tissues (55) provides additional opportunities for tissue engineering with A20. Our findings also indicate that the strategy of increasing A20 expression in the graft will synergise with other clinically tested as well as emerging graft tolerance-inducing approaches. These would include rapamycin and abatacept (4, 6, 56), as well as islet expression of the regulatory T cell attractant CCL22 (57), or graft expression of CTLA4-Ig that prevents T-cell activation (58), and local expression of the anti-inflammatory agent alpha-1-antitrypsin (59).

## Methods

### Animal models

C57BL/6 and BALB/c mice were purchased from the Animal BioResource Centre (Sydney, Australia). RAG^−/−^ and CBA mice were purchased from the Australian Resources Centre (Perth, Australia). Male, inbred, B6.129S7-Rag1^tm1Mom^/J mice were purchased from Jackson Laboratories (Bar Harbor Main USA). Neonatal porcine pancreas were obtained from 2-3 day old piglets (Swine Research and Technology Centre, University of Alberta, Edmonton AB CAN). All procedures involving animals were carried out according to the guidelines established by Canadian Council on Animal Care and the Australian institutional Animal Ethics Committee guidelines.

### Mouse islet isolation and transplantation

Islets were isolated as previously described (49), and counted for islet transplantation or *in vitro* experiments using a Leica MZ9.5 stereomicroscope. Islets were transplanted under the kidney capsule of diabetic C57BL/6 littermates at a ratio of 3 donor pancreata per 1 recipient as described (50). This strain combination represents a complete MHC mismatch. Diabetes was induced by intraperitoneal injection of 180 mg/kg streptozotocin (Sigma-Aldrich) dissolved in 0.1 M citrate buffer (pH 4.2) at a concentration of 20 mg/ml. Diabetes was determined as [blood glucose] ≥16 mM on two consecutive days measured by FreeStyle Lite® glucometer and Abbott Diabetes Care test strips following tail tipping. In some experiments transplanted mice were treated with Rapamycin or anti-PC61 monocolonal antibody. Rapamycin (LC laboratories) was dissolved in vehicle solution (0.2 % carboxylethyl cellulose, 0.25 % polysorbate-80 in 0.9 % NaCl) and administered by intraperitoneal injection on the day of transplantation and everyday thereafter for 7 days.

PC61 was used for *in vivo* depletion of CD4^+^CD25^+^ T cells, mice were injected with purified rat anti-mouse CD25 IgG1 mAb (PC61; BioExpress) intravenously (IV.; 200 μg). The efficacy of CD25 depletion was confirmed by flow cytometry. Islet grafts were retrieved from recipients at indicated time points post-transplantation for analysis of islet morphology, degree of lymphocytic infiltration by histology or gene expression by RT-qPCR. Gene expression in islet grafts was calculated using the average WT ΔCt value. Islets to be used for *in vitro* studies were cultured overnight in islet overnight culture media (RPMI-1640, 20% FCS, 100 U/ml P/S, 2 mM L-Glutamine) at 37°C + 5% CO2.

### Neonatal porcine islet isolation

Donor pancreases were surgically removed from either sex neonatal piglets and islets isolated as previously described (60).

### Adoptive Transfer

RAG^−/−^ mice were pre-transplanted with an islet graft and left to rest for 14 days. Following 14 days, spleens were obtained from mice harboring long-term surviving grafts. Harvested spleens were prepared by mechanical disruption to a single cell suspensions, erythrocytes were removed by osmotic lysis with sterile red blood cell lysis solution (0.156M Ammonium chloride, 0.01M, Sodium hydrogen carbonate, 1mM EDTA) and filtered through 70 μm nylon strainer (Becton Dickinson). CD4^+^ CD8^+^ T cell subsets were isolated via magnetic separation using Pan T cell kit (Miltenyi Biotec). Effector-T-cells (CD25^−^) were isolated from T-cell populations by positive depletion of CD25^+^ cells using the CD25 MicroBead kit following manufacturer’s instructions. Magnetic separations were performed using AUTOMACS (Miltenyi Biotec), to a purity > 95 % as assessed by flow cytometric analysis. Splenocytes T cells (2×10^7^), T cells (2×10^6^) or effector T-cells (2×10^6^) were adoptively transferred via tail vein injection.

### Cell lines

Min6 cells, generated by (61) are derived from the pancreatic beta cells of transgenic mice and immortalised by tranduction with T-antigen of simian virus 40 (SV40). MIN6 cells retain the ability to secrete insulin (62, 63). Cells were maintained in Dulbecco’s modified Eagles media (DMEM) supplemented with 10 % fetal calf serum, 2 mM L-glutamine, 12.5 mM HEPES (Gibco), 0.002% β-mercaptoethanol (Sigma), 1 % (100 U/ml) penicillin/streptomycin (Gibco) and incubated at 37°C in 5 % CO_2_. Passage 31-40 was used for experiments. Higher passage number MIN6 was avoided due to insulin secretion defects reported (63). β-TC_3_ cells, derived from insulinoma cells that arise in the pancreatic beta cells of transgenic mice expressing SV40 T antigen under the control of the rat insulin II promoter (RIP) (64, 65). These cells were cultured in Roswell Park Memorial Institute (RPMI) supplemented with 10 % fetal calf serum, 4 mM L-glutamine and 100 U/ml penicillin/streptomycin (Gibco) and incubated at 37°C in 5 % CO_2_. A passage 21-40 used for experiments.

### Recombinant adenovirus mediated gene transfer

Islets and insulinoma MIN6 cells were transduced with recombinant adenovirus (rAd.) to over-express GFP (rAd.GFP) or A20 (rAd.A20) as described previously (47, 49). The rAD-GFP construct was a kind gift from Beth Israel Harvard Medical School, Boston, MA, USA. The rAd vector expressing A20 was a gift from Dr. V. Dixit (Genentech, Inc., CA, USA). For islet gene transduction, islets were inoculated with virus at the stated multiplicity of infection, and incubated for 1.5 h at 37°C in 0.5 ml serum free RPMI-1640 medium (Gibco). Islets were then ready for further culture or transplantation. MIN6 cells were plated at a density of ~1×106/well in 6-well tissue culture plates (Corning CoStar) and inoculated with virus at the optimal MOI of 100:1 in DMEM (Gibco). After 1.5 h, cells were replenished with DMEM 10% FCS and cultured a further 24 h before use. Adenovirus was propagated by infecting HEK293 cells in six T175 vented flasks (Coring CoStar). Cells were lysed and adenoviruses were extracted using Aenopure kit according to instructions provided (PureSyn Inc.). The purified virus was titrated and quantified in HEK293 cells using the Adeno-X Rapid Titer Kit (clontech) according to manufacturer’s instructions.

### Transgene expression

GFP-expression was determined by fluorescent microscopy, images were captured under a Zeiss inverted fluorecence microscope (Carl Zeiss Inc., Jena, Germany). Islets expressing GFP were made to a single cell suspension with 0.1 % trypsin and run through a CytoFLEX (Beckman) or CantoII (BD) flow cytometer to determine GFP expression level. For A20 and IκBα protein expression, primary islets or MIN6 cells were cultured for 24 h after gene transduction and lysed in RIPA buffer, ~10 μg of total protein was resolved on a 10% SDS PAGE gel and then transferred to a nitrocellulose membrane. Membranes were incubated with polyclonal anti-A20 (Abcam, UK) and IκBα (Cell Signalling Technology, USA) respectively, and followed by peroxidase labelled secondary antibodies. Signals were visualised using an ECL detection kit (Amersham Pharmacia Biotech, Australia).

### Reporter Assays

Reporter assays were carried out essentially as we have described (45, 66). For NF-κB activity, βTC3 cells were transfected with 0.3 μg of a NFκB.Luciferase reporter with 0.25 μg CMV.β-galactosidase. pcDNA vectors encoding human A20 or empty vector up to 1ug total DNA. AP-1 activity was determined by the Cignal AP1 Reporter (luc) Kit® (SABiosciences, Australia) according to the manufactures instructions. Transfection was conducted using lipofectamine 2000 (Invitrogen) as per the manufacturer’s instructions. Following transfection cells were stimulated with 200 U/ml of TNF or 200 U/ml of each IL1β, TNF and INFγ (R&D Systems). Luciferase activity was assayed in cell lysates harvested 8 h post stimulation, using a luciferase assay kit (Promega). Results were normalized to β-galactosidase or Renilla activity (Galactostar) to give relative luciferase activity. Expression plasmids and reporters were obtained and maintained as described previously (45, 66).

### Immunohistochemistry

Tissues were fixed in 10% neutral buffered formalin (Sigma-Aldrich), paraffin embedded and parallel sections (5 μm) prepared. Sections were stained with hematoxylin and eosin (H&E; Sigma-Aldrich) or for insulin, FOXP3, CD4 or CD8 followed by counterstain with hematoxylin. To stain for insulin purified rabbit anti-mouse insulin polyclonal antibody 4590 was used (Cell Signaling Technology) followed by a HRP-labelled polymer-conjugated goat anti-rabbit IgG (Dako EnVision+ System) with DAB substrate (Sigma-Aldrich) used for visualization. To stain for Foxp3 antigen retrieval was first performed using a pressure cooker (Dako Cytomation), filled with 10 mM citrate, pH 6 (Dako Cytomation) and set to 125°C with 30 s at the maximal pressure set to 10 psi. Polyclonal anti-mouse/rat Foxp3 was used for primary antibody staining of FOXP3 antigen (eBioscience) and followed by secondary biotin anti-rat with spacer to amplify the signal (Jackson ImmunoReseach Laboratories) and visualisation of the signal achieved by using Vectastain Elite ABC kit (Vector Laboratoires, CA). CD4 and CD8 staining of paraffin sections was conducted at St. Vincent Hospital, Darlinghurst, Australia clinical histology core. Images were captured using a Leica DM 4000 (Leica Microsystems).

### Immunoblot analysis

Primary islets were lysed in islet lysis buffer (50 mM Tris-HCL pH7.5, 1% Triton X, 1043 0.27 M sucrose, 1 mM EDTA, 0.1 mM EGTA, 1 mM Na3VO4, 50 mM NaF, 5 mM 1044 Na4P2O7, 0.1% β-mercaptoethanol; supplemented with EDTA-free protease inhibitor 1045 [Roche]), cell lines were lysed with radioimmunoprecipitation (RIPA) buffer with SDS, following relevant treatment with or without 200 U/ml of recombinant human TNF (R&D Systems). Protein concentration was measured using the Bradford assay (Bio-Rad) and total protein (20-25 μg) resolved on a 7 – 10% SDS PAGE gel and then transferred to a nitrocellulose membrane, Immobilon-P® (Merck Millipore). Membranes were incubated with anti-IκBα (9242), anti-phospho-IκBα (9256), anti-JNK (9252), anti phospho-JNK (9255) (Cell Signaling Technology); or anti-beta-actin (AC15) (Sigma-Aldrich), followed by horseradish peroxidase (HRP)-labelled secondary antibody goat-anti mouse IgG Fc (Pierce Antibodies) or donkey-anti-rabbit IgG (GE Life Sciences). HRP conjugates bound to antigen were detected and visualized by using an ECL detection kit (GE Life Sciences).

### Real Time quantitative PCR

Mouse islets or neonatal porcine were isolated and placed into 12-well non-tissue culture-treated plates (150-200 islets/well; Fisher Scientific). Following an overnight culture cells were treated with 200 U/ml recombinant human TNF or of each IL1β, TNF and INFγ (R&D Systems). In some experiments cells were also pre-treated with pharmacological inhibitors, pyrrolidine dithiocarbamate (PDTC) and SP600125 (Sigma-Aldrich). Inhibitors were added at listed concentrations and incubated with cells at 37°C for 1.5 h prior to cytokine stimulation or islet transplantation. Total RNA was extracted using the RNeasy Plus Mini Kit (Qiagen) and reverse transcribed using Quantitect Reverse Transcription Kit (Qiagen). Primers were designed using Primer3 (67) with sequences obtained from GenBank and synthesized by Sigma Aldrich (Supplemental Table 1 and 2). PCR reactions were performed on the LightCycler® 480 Real Time PCR System (Roche) using the FastStart SYBR Green Master Mix (Roche). Cyclophilin (CPH2) and ACTB were used as housekeeping genes and data analysed using the 2ΔΔCT method. Initial denaturation was =performed at 95° C for 10 sec, followed by a three-step cycle consisting of 95° C for 15 sec =(4.8° C/s, denaturation), 63° C for 30 sec (2.5° C/sec, annealing), and 72° C for 30 sec (4.8° C/s, elongation). A melt-curve was performed after finalization of 45 cycles at 95° C for 2 min, 40° C for 3 min and gradual increase to 95° C with 25 acquisitions/° C.

### Flowcytometry

Flow cytometry Flow cytometric staining was performed as described (27). Data were acquired with CytoFLEX (Beckman) or CantoII (BD) flow cytometer and analysed using FlowJo software (Tree Star).

### Statistics

For gene expression studies statistical analysis was performed using the Student T-test. For allograft survival, day of rejection was plotted as Kaplan Meier curves and analyzed using the Logrank test. Tests were conducted on Prism (v5) software (GraphPad Software).

## Author contributions

Mouse islet isolation, transplantation experiments, adoptive transfer experiments and post-transplant graft analysis: N.W.Z., S.N.W. and S.T.G. Molecular studies for cell lines, islets and neonatal porcine islets and data analysis: N.W.Z. and S.T.G. Neonatal porcine isolation and transplantation: K.L.S., S.T.G. and G.S.K. N.W.Z. and S.T.G. co-wrote the manuscript. All authors read and approved the manuscript. S.T.G. conceived and designed the study.

## Acknowledgments

We thank Dr Jeanette Villanueva (Victor Chang Cardiac Research Institute, Sydney Australia) for technical advice regarding the administration and use of PC61 mAb and Dr Bernice Tan (Garvan Institute of Medical Research, Sydney Australia) for technical advice regarding NF-κB reporter assays and transfection assays with MIN6 cells. We thank Professor Goodnow (Garvan Institute of Medical Research, Sydney Australia) for providing ENU-generated mice harbouring the I325N A20 mutation.

## Supplementary data

**Supplemental figure 1.**
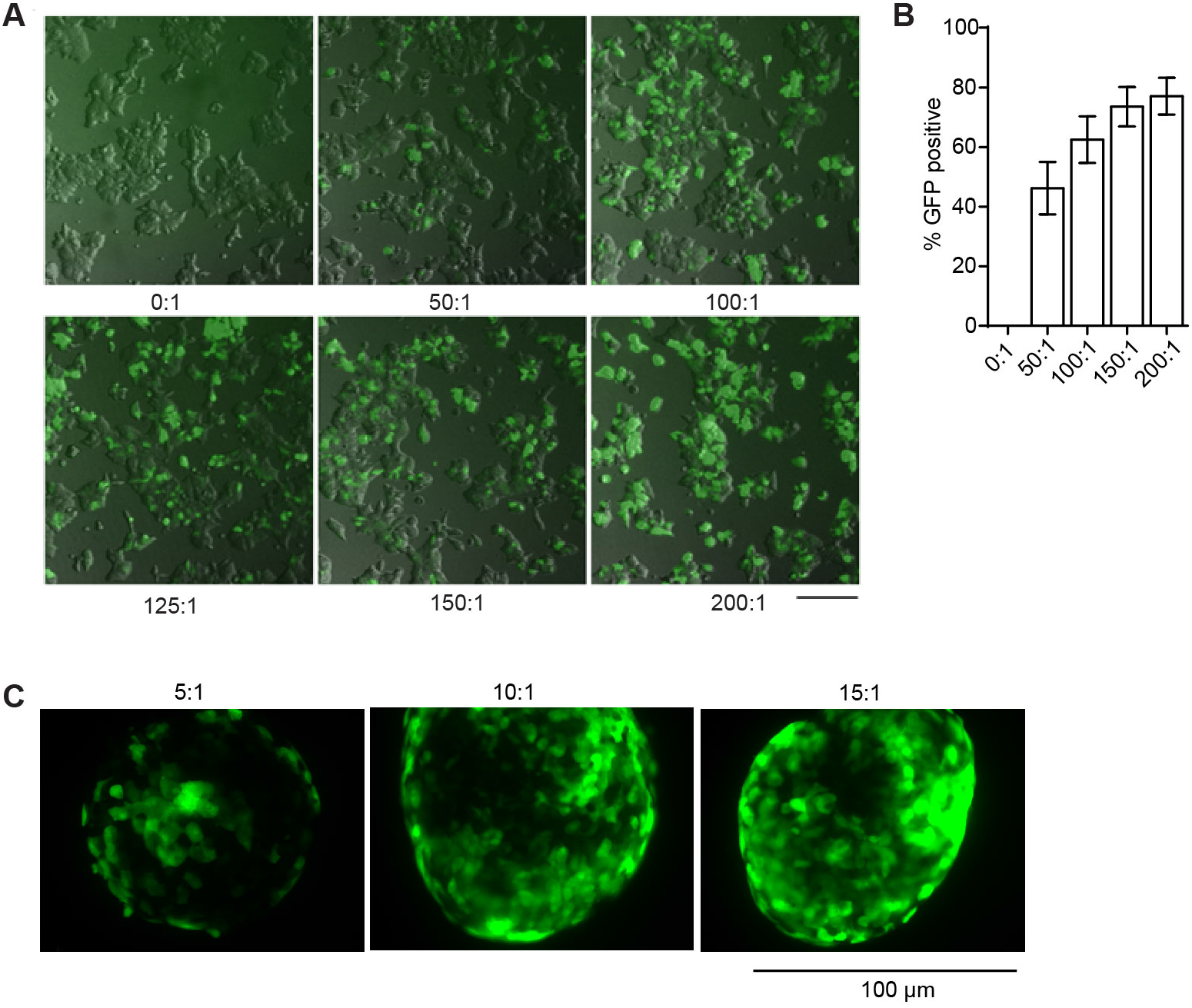
Recombinant adenovirus (rAd) transduction of MIN6 cells and mouse islets. **(A)** Representative fluorescent microscopic image of a MIN6 cell line transduced with rAd.GFP at a multiplicity of infection of 50:1, 100:1, 120:1, 150:1, 200:1 or left non-infected (NI, 0:1) after 48 h of culture (scale bar = 40 μm) and **(B)** percent of GFP positive beta cells was quantified using flow cytometry. **(C)** Representative fluorescent microscopic image of isolated primary mouse islets transduced with rAd.GFP at a multiplicity of infection of 5:1, 10:1 or 15:1 and cultured for 48 hours (scale bar = 100 μm).

**Supplemental Figure 2.**
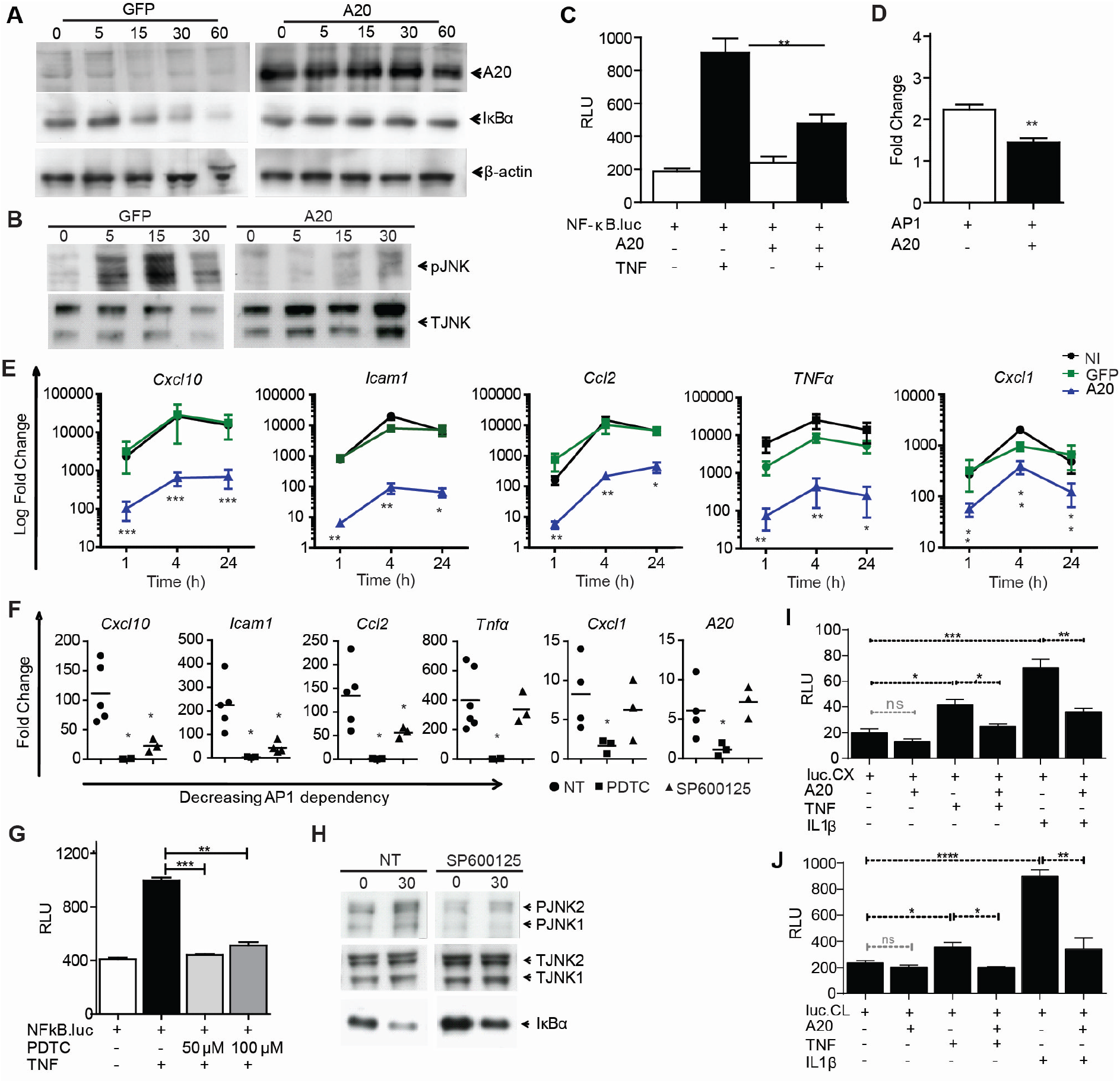
A20 inhibits TNF-induced inflammatory signalling in beta cells. **A)** Representative immunoblot of lysates from MIN6 beta cells transduced with recombinant adenovirus encoding GFP or A20 (MOI 100:1) and stimulated with TNFα (200 U/ml) for the indicated times, probed with antibodies to the indicated proteins. **(C, D)** βTC3 cells co-transfected with a NF-κB.luciferase reporter (C) or an AP-1.luciferase reporter (D) and a CMV.βgal expression construct ± PCDNA3.1 encoding A20, then stimulated with 200 U/ml TNF for 8 h or left untreated. RLU = Relative light units (Luciferase/βgal). **(E)** GFP or A20 transduced MIN6 cells or non-infected (NI) cells treated with 200U/ml of each TNF, IL-1β and IFNγ for 1, 4 and 24 h. mRNA levels were assessed by RTqPCR. **(F)** To confirm a role for NF-κB and JNK signalling MIN6 cells were pre-treated with 50 μM PDTC (to block NF-κB) or SP600125 (to block JNK) for 1 h then stimulated with 200U/ml TNF for 4 h and mRNA levels assessed by RTqPCR. **(G, H)** MIN6 cells co-transfected with a NF-κB.luciferase reporter and a CMV.βgal expression construct. Following transfection cells were pre-treated for 1 h with 0, 50 or 100 μM pyrrolidine dithiocarbamate (PDTC) and stimulated with 200 U/ml TNF for 8 h or left untreated. **(C)** Representative immunoblot of phosphorylated JNK (pJNK) and total JNK (TJNK) in MIN6 cells pre-treated with 25 uM SP600125 for 1 h or left untreated then stimulated with 200 U/ml TNFα for 30 min. **(I, J)** βTC3 cells cotransfected with a (I) CXCL10.luciferase or (J) CCL2.luciferase promoter reporter and a CMV.βgal expression construct ± PCDNA3.1 encoding A20, then stimulated with 200 U/ml TNFα or IL-1β for 8 h or left untreated. Error bars ± SEM and *P* values represent Student’s *t*-test unless otherwise stated, * *P* < 0.05; \*\**P* < 0.01; \*\*\**P* < 0.001; \*\*\*\**P* < 0.0001.

**Supplemental Figure 3.**
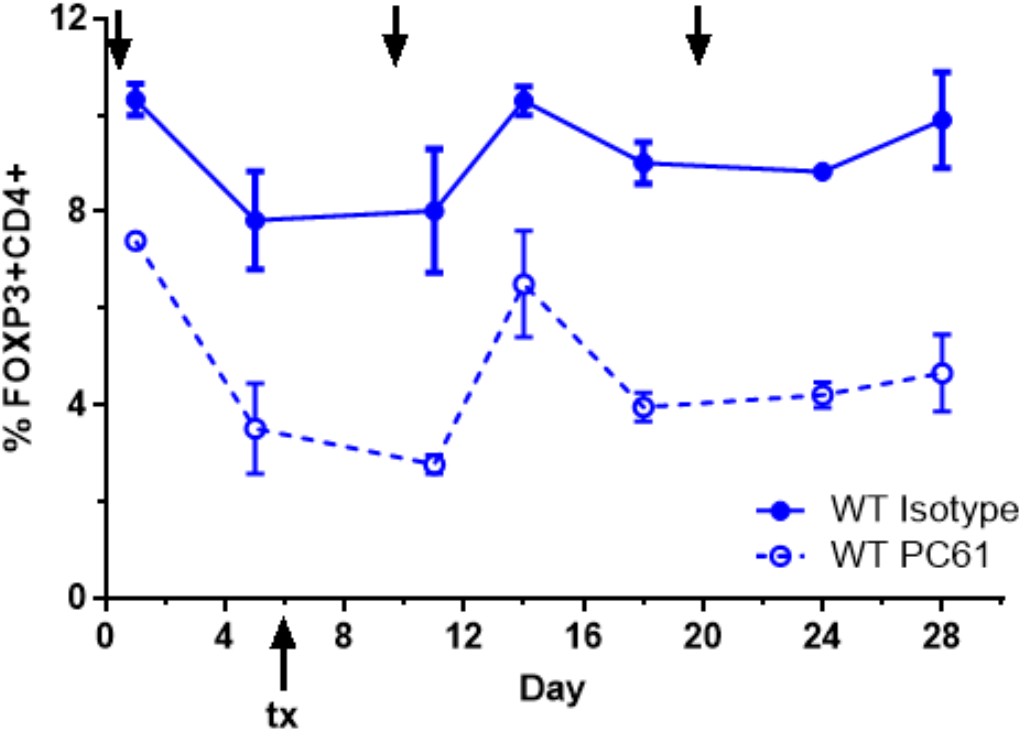
PC61 mAb treatment of wild-type mice. C57/BL6 recipients given repeated doses of αCD25 PC61, or an isotype-control and percentage of FOXP3^+^CD4^+^ T cells assessed in the blood by flow cytometry.

**Supplemental Figure 4.**
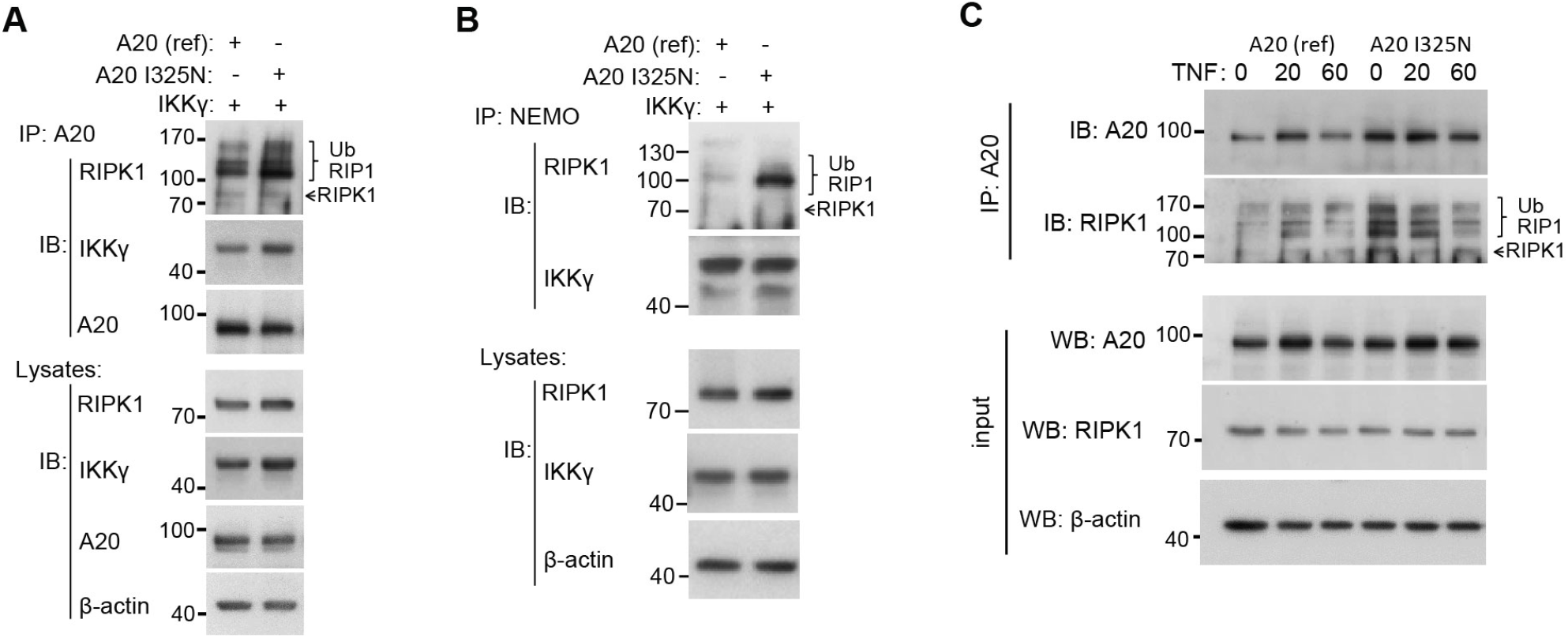
**(A, B)** Representative immunoblot (IB) of (A) A20 or (B) NEMO (IKKy) immunoprecipitated (IP) lysates from βTC3 cells transfected with reference A20 or A20 I325N and IKKy with corresponding whole-cell lysates. **(C)** Representative immunoblot of A20 immunoprecipitated (IP) lysates from βTC3 cells transfected with reference A20 or A20 I325N and stimulated with TNF for the indicated times.

**Supplemental figure 5.**
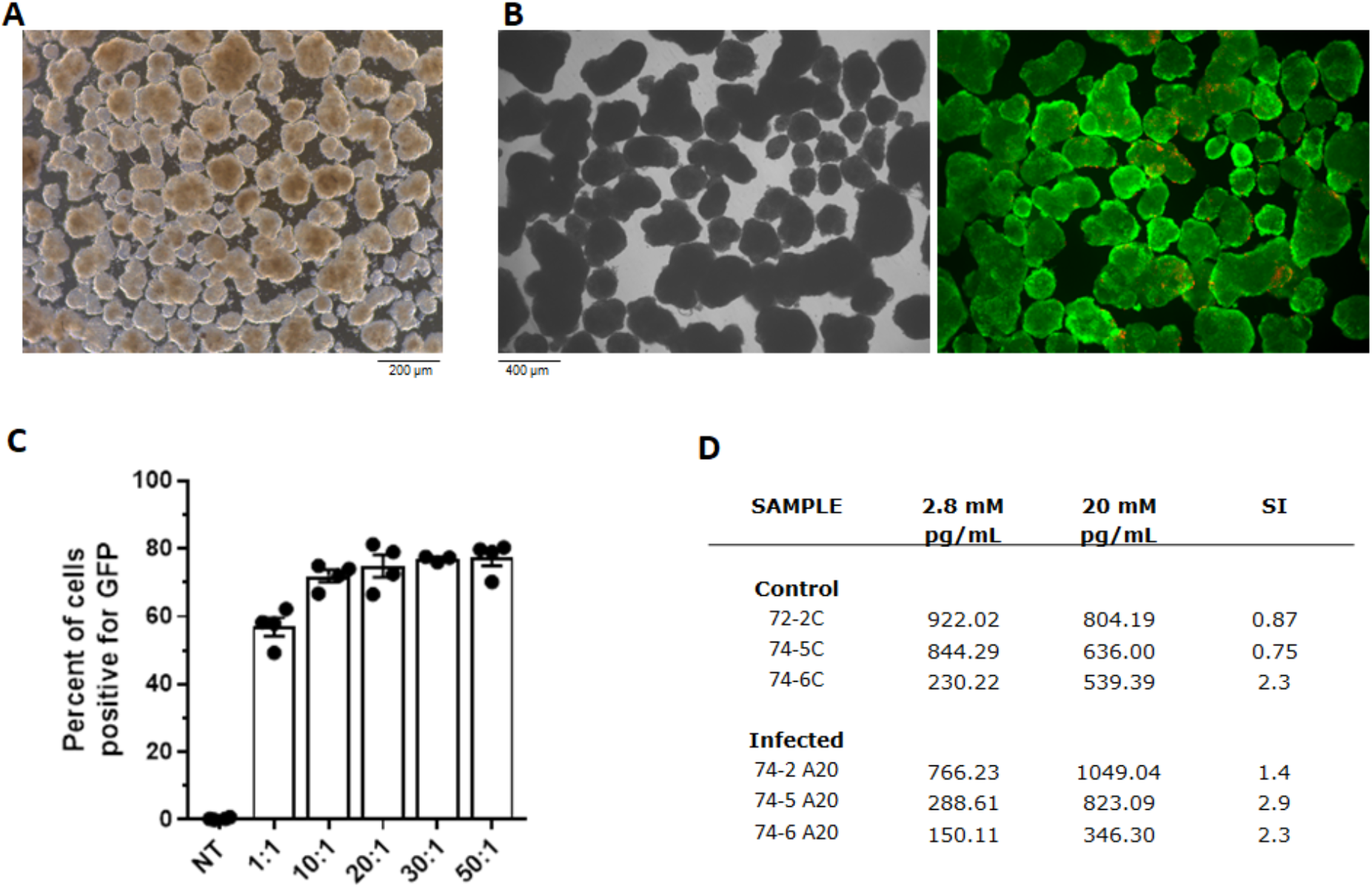
A20 expression does not impact the function of neonatal porcine islet like clusters. **(A)** Representative light microscope image and **(B)** viability stain (green = live; red = dead) of isolated neonatal porcine islet like clusters (NPIs) prior to transduction. **(C)** Isolated NPIs were transduced with an adenovirus encoding for GFP at multiplicity of infections indicated, or left untreated (NT). Forty eight hours post transduction, NPIs were digested to single cells and the percent of GFP infected cells determined by flow cytometry. **(D)** Insulin levels in supernatants of GFP or A20 transduced NPIs, incubated in 2.8 or 20 mM glucose.

**Supplemental Table 1.**
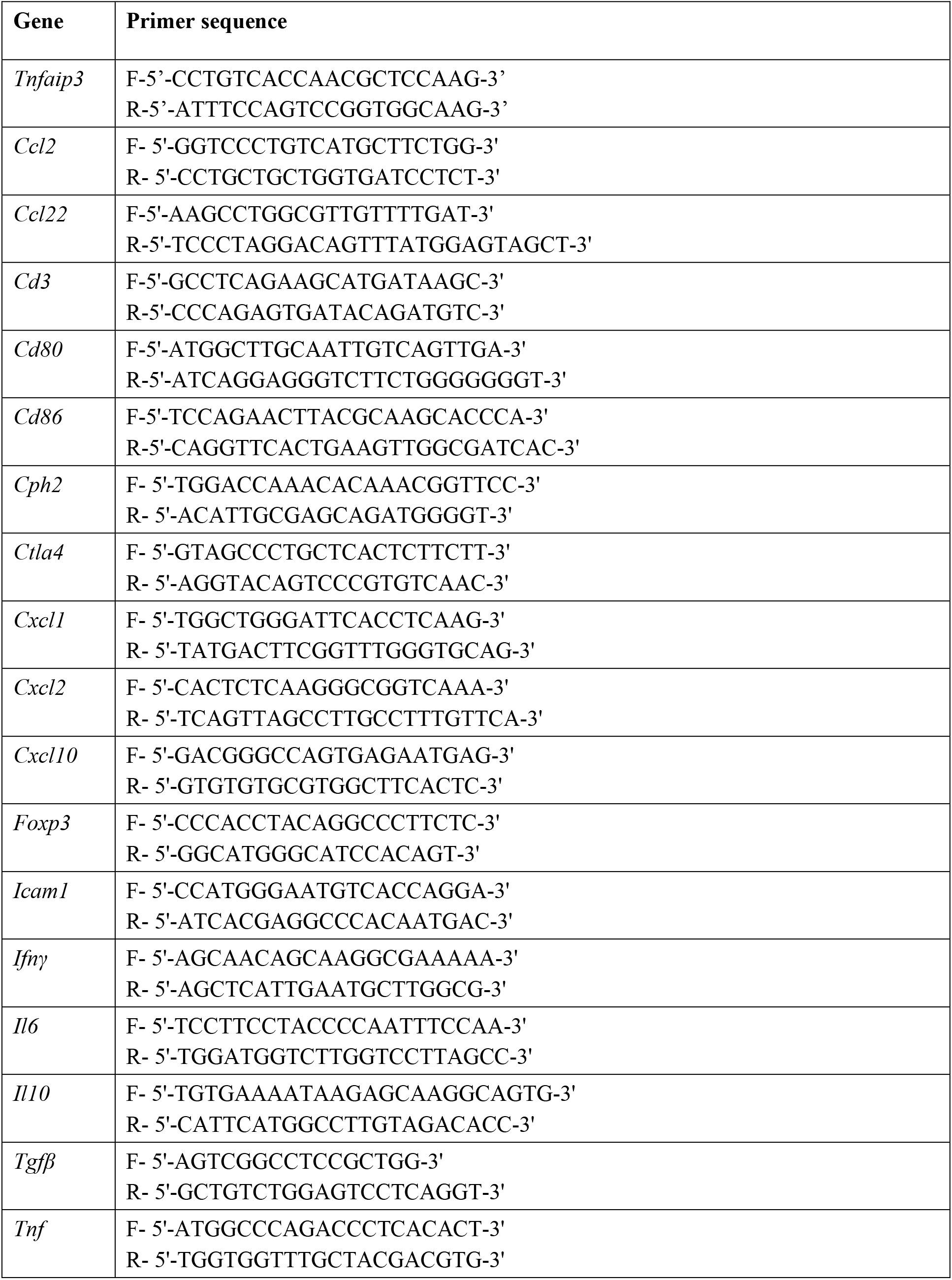
Mouse primers used for qRT-PCR analysis.

**Supplemental Table 2.**
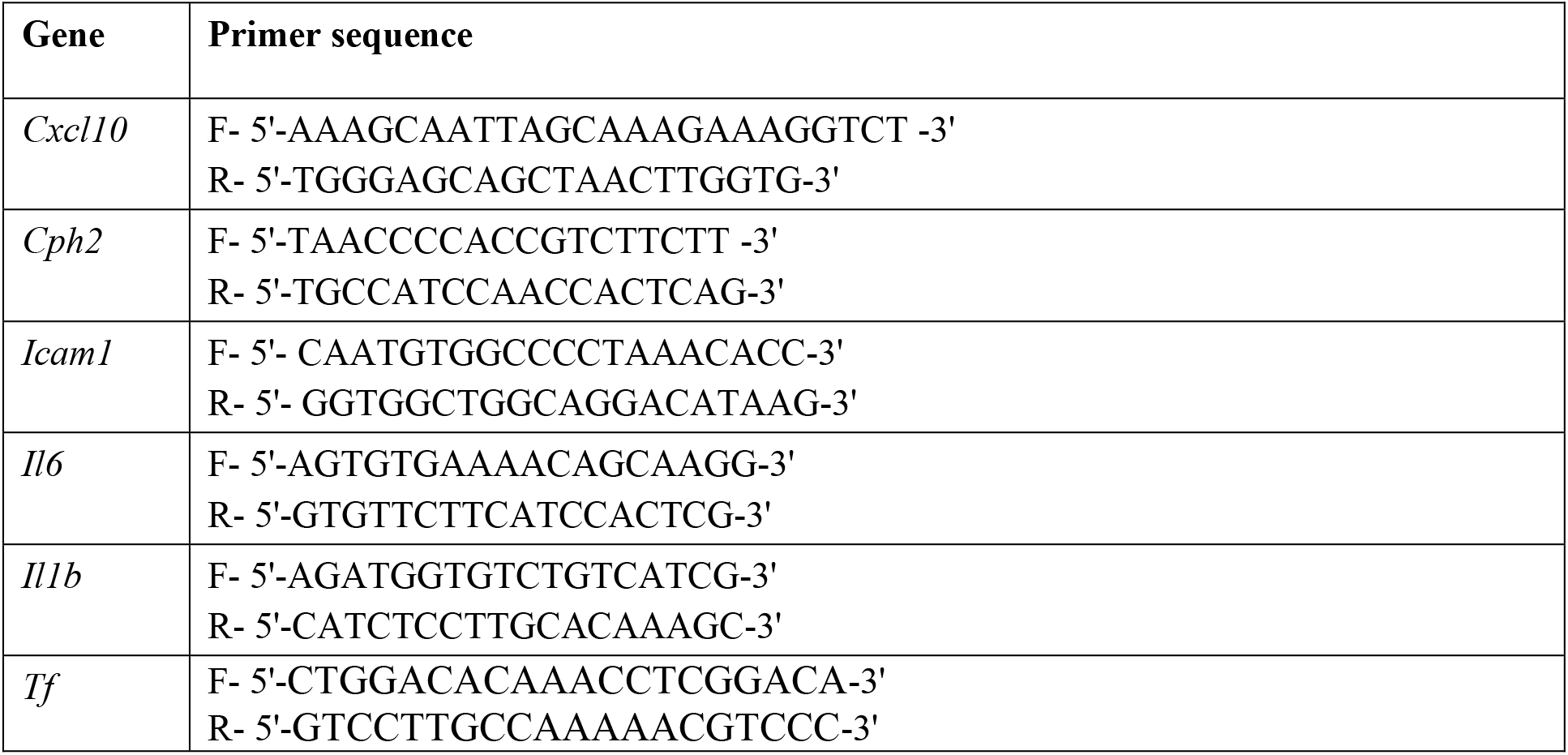
Procine primers used for qRT-PCR analysis.

